# Immunoglobulin gene conversion identification and analysis

**DOI:** 10.1101/828434

**Authors:** Stefano R. Bonissone

**Affiliations:** Digital Proteomics

## Abstract

Immunoglobulins are highly diverse, diverging from their originating germline genes driven primarily by somatic recombination and hypermutation. However, somatic gene conversion is a strong driver of immunoglobulin diversity in some species, including rabbits and chickens. It is considerably harder to detect by sequence analysis than point mutations, and currently no dedicated tools exist for identifying these events. We present `GECCO`, the first dedicated gene conversion identification tool for immunoglobulins based on modified, simultaneous, pairwise alignments to host and donor references. We benchmark our approach on simulated repertoires and find `GECCO` has high recall, low false positive rate, and is insensitive to somatic mutations. We apply this new approach to characterize gene conversion events at the repertoire level in hyper-immunized rabbits, to show patterns of donor V gene preferences and donor tract length distributions. The dedicated gene conversion identification method we present allows for the characterization of a new feature of antibody repertoires that has not been possible thus far. `GECCO` will benefit future studies to explore the prevalence of immunoglobulin gene conversion in additional species.

## 1 Introduction

Immunoglobulins display exquisite diversity as part of mounting an adaptive response to exposed pathogens. Few, if any, somatic processes induce such sequence level diversity as the immunoglobulin rearrangement and somatic hypermutation (SHM) mechanisms. It is due to these processes that novel antibody receptors can be created against newly exposed pathogens, enabling an individuals’ survival from infections. The VDJ recombination process is the first event of diversification within B cell development, mediated by RAG1/RAG2 [13]. The high level of mutation events that occur in a rearranged B cell receptor are primarily attributed to activation-induced cytidine deaminase (AID) [20].

While VDJ recombination and AID mediated SHM drive the diversity in the most studied model organisms for immunoglobulins: humans and mice, this is not the case for all animals. Another process, gene conversion (GC), is prominent in rabbits [18] and used extensively in chickens due to the avians containing only a single functional V and J gene and a large collection of V pseudogenes [1]. Furthermore, it has been shown that gene conversion is likely also mediated by AID in rabbits [30] and chickens [2].

As immunoglobulin repertoire sequencing (Rep-seq) experiments have become more commonplace, the ability to characterize features associated with immunoglobulin receptor sequences has become paramount. Determining the germline origin of a particular rearranged and mutated antibody is the first step in any downstream sequence analysis for antibody repertoire data. Being such an important preliminary step, VDJ labeling has been extensively studied and many tools exist for this task [11, 28, 7, 21, 25, 3, 12, 29, 31, 6]. By inferring the germline V, D, and J genes used, the clone (typically defined by the sequence in the CDR3/junction region) is used as a molecular fingerprint for a rearrangement event. Additionally, the inferred reference V gene is then used to determine point mutations by pairwise alignment between the read and inferred V germline gene reference. Several pipelines for analyzing Ig-seq data have been developed to perform these steps [27, 5, 26]. However, since most studies have focused on human repertoires (to study disease response or autoimmunity), or mouse (to study basic immunology and discover antibodies), gene conversion has largely been ignored. Since GC is thought to be rare in these species, no tools have been developed for the identification of gene conversion events in large scale high-throughput immunoglobulin sequencing. To date, the examples of gene conversion studies in literature rely on a tool, GENECONV [23], that was designed for low throughput analysis of gene conversion within single genes. Other studies use visual inspection in a multiple sequence alignment to confirm their suspicion of gene conversion events. Lavinder et al., [18] recreated an internal version of GENECONV for their study, while Winstead et al., [30] used visual inspection, and Duvvuri et al., [9] used both GENECONV and visual inspection. Visual inspection is low throughput and unable to systematically characterize repertoire level patterns of gene conversion, in addition to not providing a formal framework for their identification. Furthermore, we show here that the GENECONV approach lacks sensitivity to a wide range of possible gene conversion events and produces far more false positives than our proposed approach.

In this manuscript we present a novel algorithm to tackle the problem of identifying gene conversion in immunoglobulin sequences, and an implementing tool called GEne Conversion Classification of immunOglobulins (`GECCO`). This tool will enable a new category of analysis for immunosequencing datasets, now being able to accurately model immunoglobulin diversity and explore the role gene conversion plays in the adaptive immune system.

## 2 Methods

### 2.1 Immunoglobulin VDJ classification

The problem of labeling an antibody sequence by its constituent V, D, and J genes, i.e., VDJ classification, has been tackled previously in the literature [21, 28, 12, 31, 6]. Briefly, it can be described as the following: given reference gene-segment sets *𝒱, 𝒟, 𝒥*, and a read *r*, return the labels *v* ∈ 𝒱, *d* ∈ 𝒟, and *j* ∈ 𝒥 that best explain *r*. Despite this simply described problem, creating an accurate model for VDJ classification is a difficult task. As a result, this classification can be error prone, particularly for the typically more diverse heavy chain. Our approach to gene conversion identification is predicated on having a highly likely V gene identified as the host gene and another as the donor gene. While ensuring high accuracy on the top ranked V gene is typically difficult, if considering the top three, this problem becomes considerably simpler. Our approach uses the top *m* sequences using them all as candidate host and donors, and as such, requires having performed VDJ classification, if not provided, `GECCO` will run VDJ classification to obtain a ranked list of V gene candidates.

`GECCO` performs this classification task much as [29, 3, 31], by first aligning V genes, removing aligned sequence to the reference, then aligning J genes and removing the matching substring, and finally aligning to the D genes. This iterative alignment classification works well enough for our purpose, however, since large gene conversion events can make the prediction of the host V reference more error-prone, we must check the top *m* not only for the donor, but also host reference.

### 2.2 Gene conversion identification

Immunoglobulin gene conversion (GC) events are common in some organisms, such as rabbit [18] and chicken [18], while there is some evidence that it rarely occurs in human [8, 9]. Despite the fact that studies have analyzed GC events from limited sequencing data, there does not exist a tool for identifying gene conversio in immunoglobulin sequences. Lavinder et al., 2014 [18] approach analyzing GC using a variation of the GENECONV permutation test. Duvvuri et al., 2012 [9] use two approaches: GENECONV [23] and visual assessment.

Gene conversion events appear in sequence analysis as a donor substring (termed tract) inserting, and deleting, a portion of the host nucleotide sequence. There exist tools for identifying gene conversion events at the genomic level [24, 19, 14] (i.e., non-somatic), but assumptions make their application to immunoglobulins difficult, particularly, collapsing sequences into consensus sequences prior to comparison [19, 14]. While general tools for split read mapping could be modified [10] or a seed-extension strategy could be employed [16], we take a direct approach for immunoglobulin somatic gene conversion.

We present the problem of gene conversion identification as one of identifying substring insertions from a donor germline V gene-segment *v*_*d*_ to a host germline V gene-segment *v*_*h*_. Given a query *r*, and set of reference V genes 𝒱, identify the ordered set of substrings, e.g., 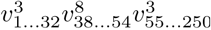, that maximize the coverage over the query *r* while penalizing each reference gene used, *v*^3^ and *v*^8^ in the example. Our approach to gene conversion identification can be seen as an extension of VDJ classification, where two references are allowed instead of one. The use of two references poses a problem of overfitting, much like the problem of choosing the correct number of clusters *k* when performing unsupervised clustering. In a similar vein, we compare the gene conversion model score (similar to *k* = 2 clustering) against the single host V reference score (akin to *k* = 1 clustering), to determine which is a better fit given the additional parameters.

The venerable GENECONV [23] program is the only cited tool in the literature that has been used to analyze GC events in immunoglobulins. GENECONV takes as input a multiple sequence alignment, removing any monomorphic columns (i.e., columns containing an identical nucleotide across all rows), identifying pairs of identical substrings, and using a permutation test to determine if each pair of tracts are significant. However, this permutation based approach has certain pitfalls. Given a large alignment, permuting the columns 10,000 times can be time consuming if performed for millions of multiple alignments - even when only considering polymorphic columns. Furthermore, the approach is sensitive to which sequences are contained in the multiple alignment. Meaning, given *n* sequences, the resulting fragment could likely differ if an alignment with one additional sequence was provided instead; this sensitivity to the input sequences is particularly problematic. Furthermore, GENECONV as used for identifying tracts in immunoglobulin sequences requires the correct donor template sequence to be present in the multiple alignment. If too many sequences are added to the multiple alignment reduced sensitivity, as well as a computational slowdown occur. More fundamental, however, is that a multiple sequence alignment is not well suited as it attempts to align all sequences to one another. Unlike the approach to GENECONV, the true task is to align each reference to the read, and not to each other which can introduce artifacts.

#### 2.2.1 GC identification by GENECONV

Using a similar approach to [18], is to use GENECONV to perform the identification of a segment shared between the read and a donor reference sequence. The input required to GENECONV is a multiple sequence alignment containing the read *r*, host reference *v*_*h*_, and putative donor reference *v*_*d*_. The host reference is selected as the best matching V reference by the VDJ classifier used, e.g., IgBlast. The putative donor references are the subsequent *m*−1 V references identified (i.e., references ranked 2 to *m*). With a read *r*, host reference sequence *v*_*h*_, and donor sequence *v*_*d*_, then the three can be aligned in a multiple alignment *A*. At which point, only polymorphic columns (i.e, columns that have at least one difference across all rows) are retained, and a multiple alignment with fewer columns *A*′ is obtained. The longest tract of the donor segment is identified in *A*′, and the tract is then expanded back into the full alignment of *A* to obtain the full donor tract sequence. The donor tract length in *A*′ is used as the query test statistic for the permutation test, which then permutes the columns of *A*′ 10,000 times, recomputing the longest common donor tract length each iteration. The query test statistic is then compared against the permuted, null distribution to obtain an empirical p-value for the original query. This algorithm described is sketched in Algorithm 1.

#### 2.2.2 Gene conversion identification by mirrored alignment

A direct approach to solving the gene conversion identification problem is to use dynamic programming and combine pairwise alignments between the read sequence and two different references. Specifically, for the query *r*, and references *v*^1^ and *v*^2^, *align*(*r, v*^1^) and *align*(*r, v*^2^) are computed simultaneously, allowing jumps between them, representing gene conversion events; i.e., *v*^1^ and *v*^2^ represent putative host and donor references. More specifically, two alignment matrices, *A*_1_ and *A*_2_ are created, for aligning the query to the two references. The recurrences for the two matrices are similar:

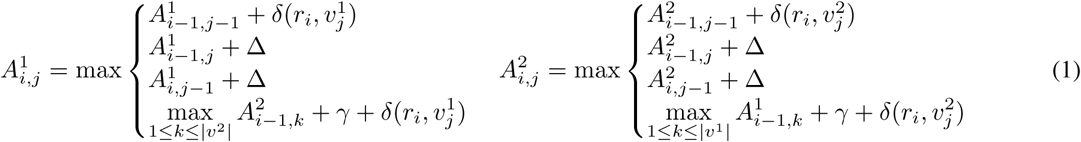

The recurrences for each matrix are the same as the Needleman-Wunch alignment, with the additional allowance of a transition from *A*^1^ to *A*^2^ for a penalty of *γ*. This transition allows jumps from anywhere between columns 1 and |*v*^1^| or |*v*^2^| (shown in Figure S1), which can be performed more efficiently by using an additional matrix to keep track of the maximum column jump for each row. Backtracking starts at the maximum scoring final position of the two matrices, max 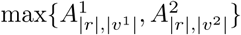, following the standard backtracking procedure, allowing for jumps between the matrices. In addition to backtracking providing an alignment between the read *r*, and the composite references of *v*^1^ and *v*^2^, additionally a trace string *b* is returned, noting which matrix the alignment position originates from: *A*^1^ or *A*^2^. The use of equation 1 to call a gene conversion event is further described in Algorithm 2.

### 2.3 Statistical assessment of GC alignments

The alignment procedure obtained using equation 1, *AlignGC*, will always produce an alignment, trace, and score. However, merely because an alignment jumps between the host germline *v*_*h*_ and the donor germline *v*_*d*_ does not ensure that this will be called as a gene conversion event. Instead we assess the statistical significance of a gene conversion event by comparing the result of *AlignGC*(*v*_*h*_, *v*_*d*_, *r*) to that of traditional pairwise alignment to only the host reference, *Align*(*v*_*h*_, *r*). By rescoring each of these alignments, we can frame them as a likelihood ratio test and exploit the ability to compute a p-value using the *χ*^2^ distribution.

The alignments produced by *AlignGC* and *Align* are then rescored using a simple probabilistic model that takes into account mutations using a match/mismatch penalty matrix *δ*(·,·), and a probability of initiating a gene conversion event, *γ*. Such a likelihood model takes parameters *L*(*A*|*δ, γ*) to provide an estimate for alignment *A*. In the case of standard pairwise alignment, *Align*, we could use *AlignGC* with the gene conversion parameter *γ* = −∞, ensuring that only alignment of the host reference *v*_*h*_ and read *r* are enforced. This model nesting suggests that a likelihood ratio test is appropriate. Specifically, the likelihood function can be computed for the fully parameterized model, *L*(*A*|*δ, γ*),and compared to a partially parameterized model *L*(*A*|*δ*). Then, the test can be set up as:

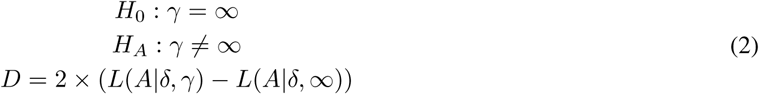

The likelihood ratio test above can be assessed for significance using the *χ*^2^ distribution with a number of degrees of freedom equal to the difference in number of parameters in models used between *H*_0_ and *H*_*A*_; in this case 1. The test statistic, *D*, is thusly compared to the threshold for type I errors, *χ*^2^(*D*, 1) ≤ *α*. This provides a method for controlling false positives, by comparing a non-GC model to the GC model. The test does not control the *false call* rate (FC); when a gene conversion call is made correctly, but the parameters of the prediction are incorrect. This occurs when a threshold on the similarity of the predicted GC tract interval is used to determine a true call. The likelihood ratio test alone only compares the GC model against the mutation only model for significance. In other words, testing if the differences between the sequence and reference can be explained only with mutations, or if additional parameters (i.e., modelling gene conversion) are necessary.

False call errors can be categorized into three types that generate an incorrectly predicted tract interval: 1) the incorrect host V gene was predicted; 2) the incorrect donor V gene was predicted; and 3) the correct host and donor were predicted, but the interval is incorrect. Errors of incorrect host V gene are very difficult to correct, and are a minority of errors. Errors of donor V genes are more common, and are typically caused by limiting to the top *m* V genes. We will show that these errors can be minimized by increasing *m* and considering more donor V genes at a modest expense of compute time. Finally, most errors are from calling the incorrect interval, caused by the donor interval being too small to detect, or more often, the interval is in an indistinguishable tract of host and donor sequence.

## 3 Results

### 3.1 Datasets

#### 3.1.1 Simulated antibody datasets

In order to evaluate gene conversion identification sensitivity to known mutation loads and gene conversion tract properties, simulated antibody sequences are generated. Simulated antibody (smAb) sequences are created in a straight-forward manner. First, a host V, D, and J reference is selected uniformly at random. Furthermore, the number of nucleotides that are removed from each end of the 3’ V, 5’D, 3’D, and 5’J references are sampled from a published distribution [15]. While non-templated nucleotide composition and length are sampled from a different published distribution [4].

Once a recombined smAb is generated it can then be subjected to mutations, sampled from an internally derived position specific distribution. Gene conversion tract starting position are selected uniformly random across *v*_*h*_, while the tract lengths are sampled from a Gaussian distribution. A repertoire of smAbs can be generated with different population level properties as well, e.g., uniformly random germline usage or ensuring expanded clones are created. For the purposes of experiments in this manuscript, uniform distributions were created, unless otherwise noted, so as to exhaustively cover the gene conversion events observable in Rep-seq datasets.

#### 3.1.2 Rabbit immunosequencing dataset

Three juvenile New Zealand white rabbits were used, one for each repertoire dataset. For each immunogen, two rabbits of random sex were immunized in complete Freund’s adjuvant. Additional boosters of incomplete Freund’s adjuvant were administered at later weeks, differing schedules for each immunogen. One rabbit was an extension of an existing immunization, two additional boosts of recombinant antigen of conch carrier protein were performed at weeks 0, and 3 of the extended boost. Rabbit peripheral blood mononuclear cells (PBMCs) and bone marrow were collected during the course of immunization. Additional PBMCs were collected at weeks 1, 2, 4, 5, 6, and 7, and bone marrow collected at the terminal bleed. Two additional rabbits were boosted with recombinant protein antigens during a normal 14 week schedule (boosts at weeks 3, 6, and 10) and PBMCs collected at weeks 4, 7, 9, 11, 14, and bone marrow collected at the terminal bleeds. Serum titers were measured via ELISA for productive IgG response, and rabbit producing highest titer against the antigen at 7 weeks was selected analysis. Bleeds were performed pre- and post-boost, along with a final bleed at 14 weeks. PBMCs were isolated with Ficoll gradient by Pacific Immunology from pre-immune, pre-boost, and post-boost bleeds. Rabbit processing was performed by Pacific Immunology (NIH OLAW Assurance #:A4182-01, USDA registration #: 93-R-283).

Sequencing libraries were created using proprietary paired forward and reverse primers targeting transcripts from the IGH and IGK loci. The resulting libraries were sequenced on an Illumina MiSeq using the 2×300nt chemistry. Reads were processed and transformed into a repertoire using an internal pipeline. Briefly, preprocessing and paired-end read assembly provide full length reads covering the entire V region. Error correction proceeds using a Hamming graph approach [22], VDJ classification performed using iterative alignment, and subsequent mutation analysis and compilation into an annotated repertoire. Three heavy chain repertoires were obtained, A12, A14, and A18, and A14 will be the focus of analysis in the main text.

### 3.2 Gene conversion identification

#### 3.2.1 Evaluating putative gene conversion events

In order to assess the ability to identify gene conversion events, different criteria for correctly identifying a gene conversion event within an smAb are established. The simplest criteria is determining if the correct donor V gene ID is identified. The second criteria is comparing the identified gene conversion tract interval to that of the known donor location within the host (for simulated reads). Comparison of the true tract interval *d*_*t*_ and predicted tract interval *d*_*p*_ is performed using the Jaccard similarity:

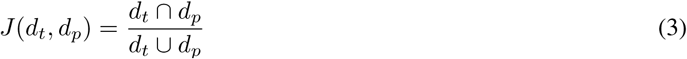

The Jaccard score ranges from [0, 1], with 1 occurring when *d*_*t*_ = *d*_*p*_, and 0 when there is no overlap between *d*_*t*_ and *d*_*p*_ intervals. A Jaccard score is computed for each predicted gene conversion event on simulated data, with a gene conversion identification method producing a distribution of higher Jaccard scores indicates better performance. Additionally, the Jaccard score can be thresholded to provide a correct or incorrect label for an event, *J* (*d*_*t*_, *d*_*p*_) ≥ 0.5 is used for correctly called tracts.

Bootstrap sampling is performed for each tract length or mutation load parameter tested, resampling 1,000 sequences from a population of 10,000 for ten iterations. Each statistic presented is the mean value over these ten iterations with 3*σ* standard deviations shown as error bars over every point, unless otherwise noted.

Any predicted donor tract that has a start or end position in the first or last 10nt of the host V gene is not considered a gene conversion event. This is to ensure that predicted donor tracts come from likely gene conversion events (i.e., two breakpoint recombination events) and not from PCR chimeras (i.e., single breakpoint recombination events).

#### 3.2.2 Results on simulated gene conversion repertoires

To test the performance of `GECCO` and GENECONV for identifying tracts of gene conversion, simulated antibody sequences were generated with different mutation loads and gene conversion tract length distributions. Prior to evaluating GC-ALIGN, additional datasets were simulated to determine a well performing parameter value for *γ* using line search, Figure S2, while keeping other alignment parameters fixed.

Figure 1 shows the performance of our GENECONV implementation and `GECCO` when using the identified host reference and *m*−1 putative donor references from VDJ classification. It also shows the best potential performance when using the host and donor references used to generate the simulated antibody. Particularly concerning is the difference between the best case scenario with known host and donor references (Figure 1a) and the donor blinded approach (Figure 1b), demonstrating that GENECONV has relatively low sensitivity for tracts shorter than 80nt, even when a small number of mutations are present. GC-ALIGN shows better performance over all ranges of donor tract lengths for the simulated rabbit repertoires, and similar superior performance is also observed in simulated human repertoires (Figure 1c, Figure 1d, and Figure S3).

**Figure 1:**
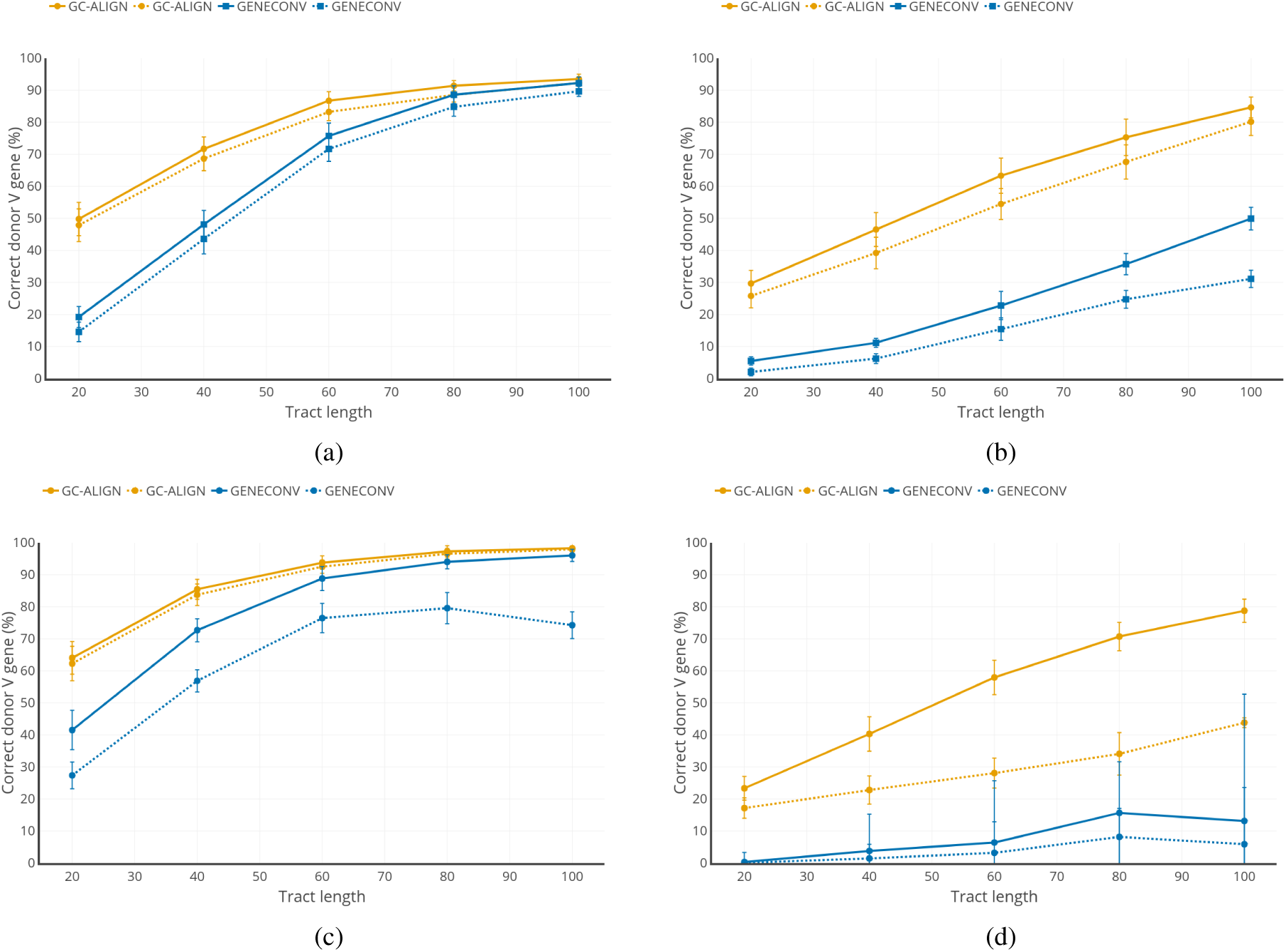
Results on simulated gene conversion data. The two types of lines show the percentage of gene conversion events called at a fixed *α*: the percentage of identified events based solely based on donor V gene ID; and the percentage of events identified by Jaccard similarity ≥ 0.5 on the predicted and true intervals. All GENECONV gene conversion events were called when the permutation p-value ≤ 0.05, and GC-ALIGN also used a p-value ≤ 0.05 cutoff. Results on simulated reads with varying levels of mean gene conversion donor tract length, each with a standard deviation of 10nt; rearranged and gene converted sequences contain no mutations. Bars signify 3*σ* deviations. Left column are results when using known host and donor V genes as input, right column are results from a completely blinded method, performing V gene identification prior to gene conversion identification. a) Simulated rabbit repertoire identified with known host. b) Simulated rabbit repertoire identified with blinded host. c) Simulated human repertoire identified with known host. d) Simulated human repertoire identified with blinded host.

In addition to identifying gene conversion events, each method predicts an interval in the host reference that originates from the donor reference. Figure 2a shows the distribution of Jaccard similarities of predicted gene-converted donor tracts compared to the true donor tract intervals. GC-ALIGN has better predicted tracts for all donor lengths. Detailed density plots of the Jaccard distributions are shown in Figure S4, and slight biases in calling boundaries are shown in Figure S4a.

**Figure 2:**
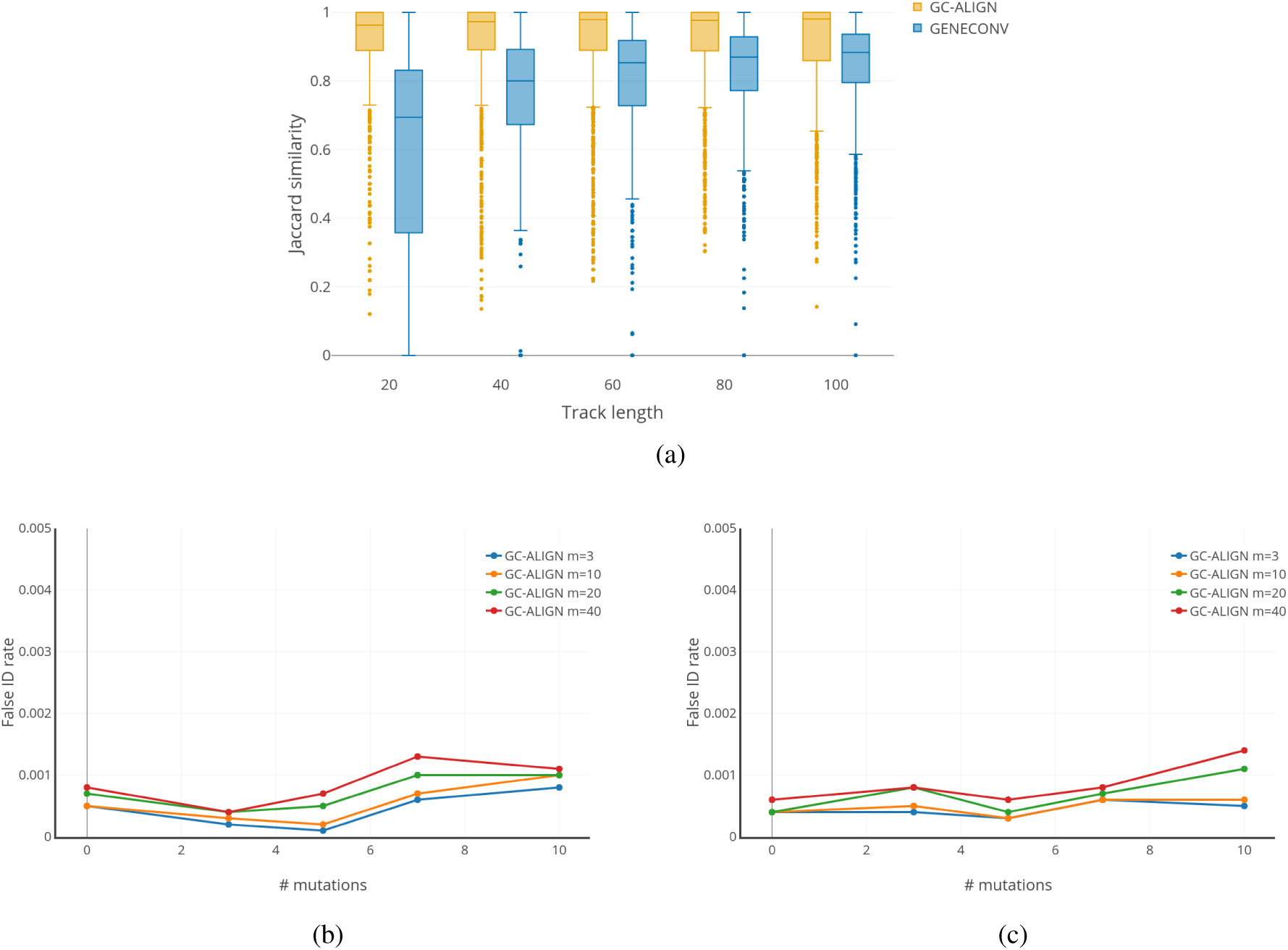
a) Jaccard similarity of identified donor sequence intervals compared to the true interval of inserted donor sequences on rabbit simulated data from Figure 1. b) and c) show false positive rates for human simulated datasets of 10,000 sequences each with a specified number of nucleotide mutations; b) uniformly distributed, and c) distributed preferentially in CDR1 and CDR2, at a 5:1 ratio compared to framework regions. The lines follow the parameter *m* (the number of references to check donors), with the largest *m* = 40 incurring the highest false positive rate.

Another consideration is that somatic hypermutations (SHMs) could reduce sensitivity of different methods. Figure 3a shows the accuracy of our GENECONV and GC-ALIGN with datasets of varying levels of mutations when provided with known host and donor references. GENECONV is sensitive to the level of SHMs, while GC-ALIGN is insensitive to mutation load. GENECONV showed considerable loss of recall as mutation load increases up to recalling only 10% of events when ten mutations are present, while GC-ALIGN shows higher recall at all levels of mutation (Figure 3b). Additionally showing that increasing the number of putative donors provided by VDJ labeling can improve recall considerably (also seen in human in Figure S3).

**Figure 3:**
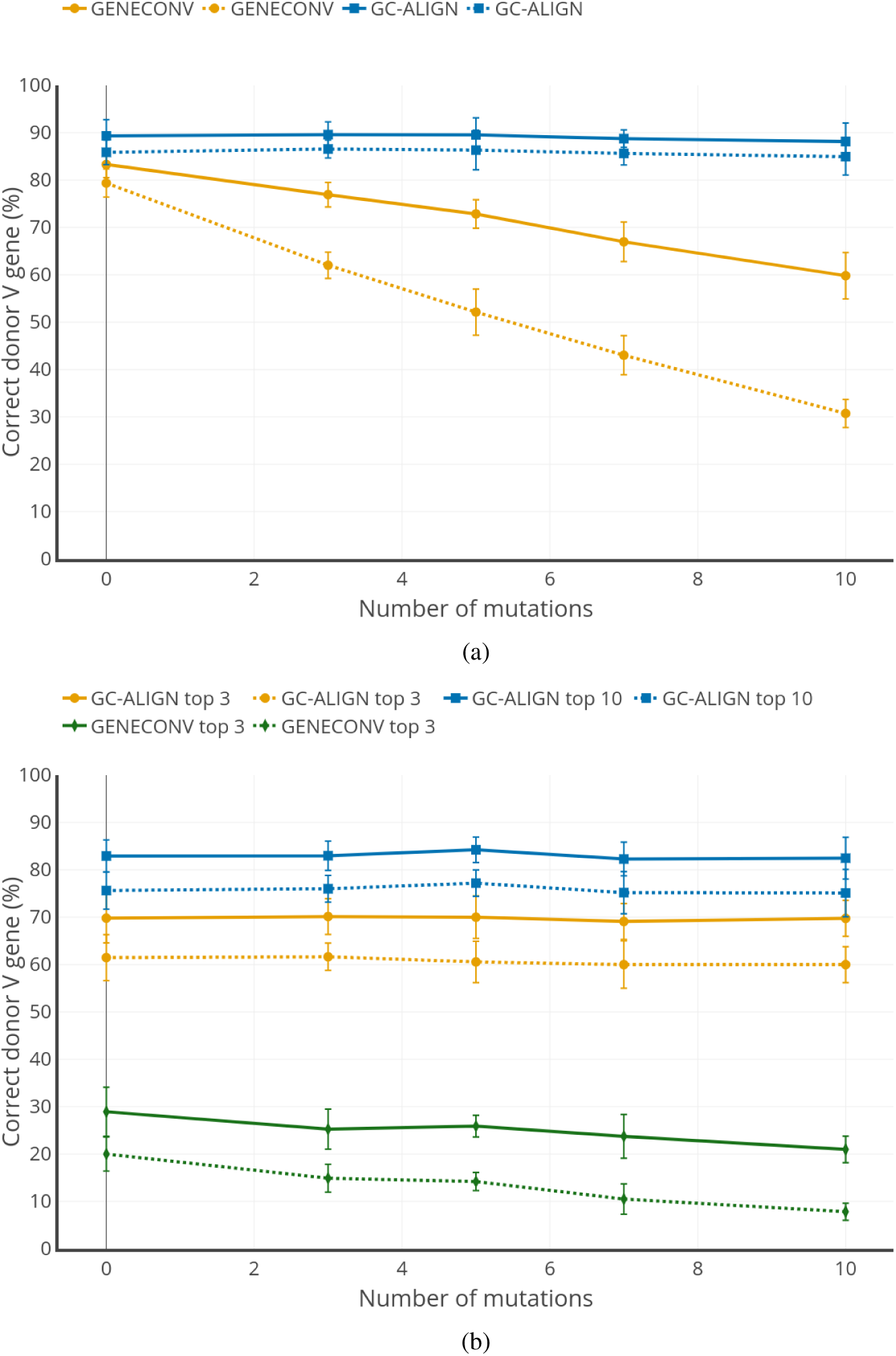
Performance of gene conversion identification at increasing numbers of nucleotide mutations with a mean donor tract length of length 70nt±10nt. Solid lines show correct gene conversion event, while the dashed line shows the correct donor interval by Jaccard similarity ≥ 0.5. a) True host and donor V genes are used as input, GC-ALIGN is not sensitive to the number of mutations in the V gene-segment, while GENECONV loses sensitivity as the number of mutations increase. b) Host and donor genes are inferred. Using the top 3 V genes has reduced sensitivity compared to using the top 10.

Furthermore, GENECONV commits more false positive errors on simulated rabbit repertoires that have been mutated, but not gene converted. Simulating 10,000 sequences with 0, 5, 10, and 20 nucleotide mutations, the GENECONV approach resulted in 250, 213, 219, and 190 false positive calls; while GC-ALIGN commited 4, 4, 4, and 25 false calls. GENECONV had a false positive rate of 1.9%, while GC-ALIGN had a rate of 0.25%, on simulated sequences with the most number of mutations tested. This low false positive rate persisted when testing simulated human repertoires with varying numbers of donor references (i.e., varying *m*) at different levels of mutation, with false positive rates between 0.001-0.002 (Figures 2b and 2c).

#### 3.2.3 Gene conversion in rabbits

Previous studies report rabbit repertoires predominantly use the VH1 gene, which is likely due to the VH1 gene’s location immediately upstream of the D gene cluster on the IGH locus. Three different VH1 allotypes have been previously characterized [17], VH1-a1 (IGHV1S69*01), VH1-a2 (IGHV1S34*01), and VH1-a3 (IGHV1S40*01). The rabbit from the A14 repertoire had a dominant host allotype IGHV1S69*01 and also expressed a minor host allotype IGHV1S34*01, suggesting it was heterzygote for the two allotypes. Rabbits from A18 and A12 utilized only the IGHV1S34*01 host allotype, but the dominant host V gene is assigned to IGHV1S7*01.

Repertoires from three different rabbits, immunized with three different protein antigens were created and processed using an internal pipeline (Table S1). The datasets A12, A14, and A18 generated 202,309; 212,955; and 162,240 unique reads, respectively. Results from A14 are presented in the main text, which contains repertoire reads from 8 different heavy chain libraries across 7 time points in PBMC and bone marrow compartments. These data were processed to obtain 96,736 unique sequences with read count of five or more, which were used for the analysis to ensure the sequences are likely error free. These sequences were analyzed for gene conversion using GC-ALIGN, and the summary of results can be seen in Figure 4. Specifically, V1S69 consisted of the vast majority of host sequences, while gene V1S44 was the most prominent donor and almost exclusively donated tracts to V1S69, Figure 4a. In total, 22,310 of the 96,736 entries were predicted to contain GC events (23.06%). The other two datasets, A12 and A18, had similar rates of GC with 23.55% and 19.99% called GC events, respectively. The distribution of gene converted donor tract lengths is shown in Figure 4b. The bulk of this distribution shows tract lengths between 50nt-80nt are the most common.

**Figure 4:**
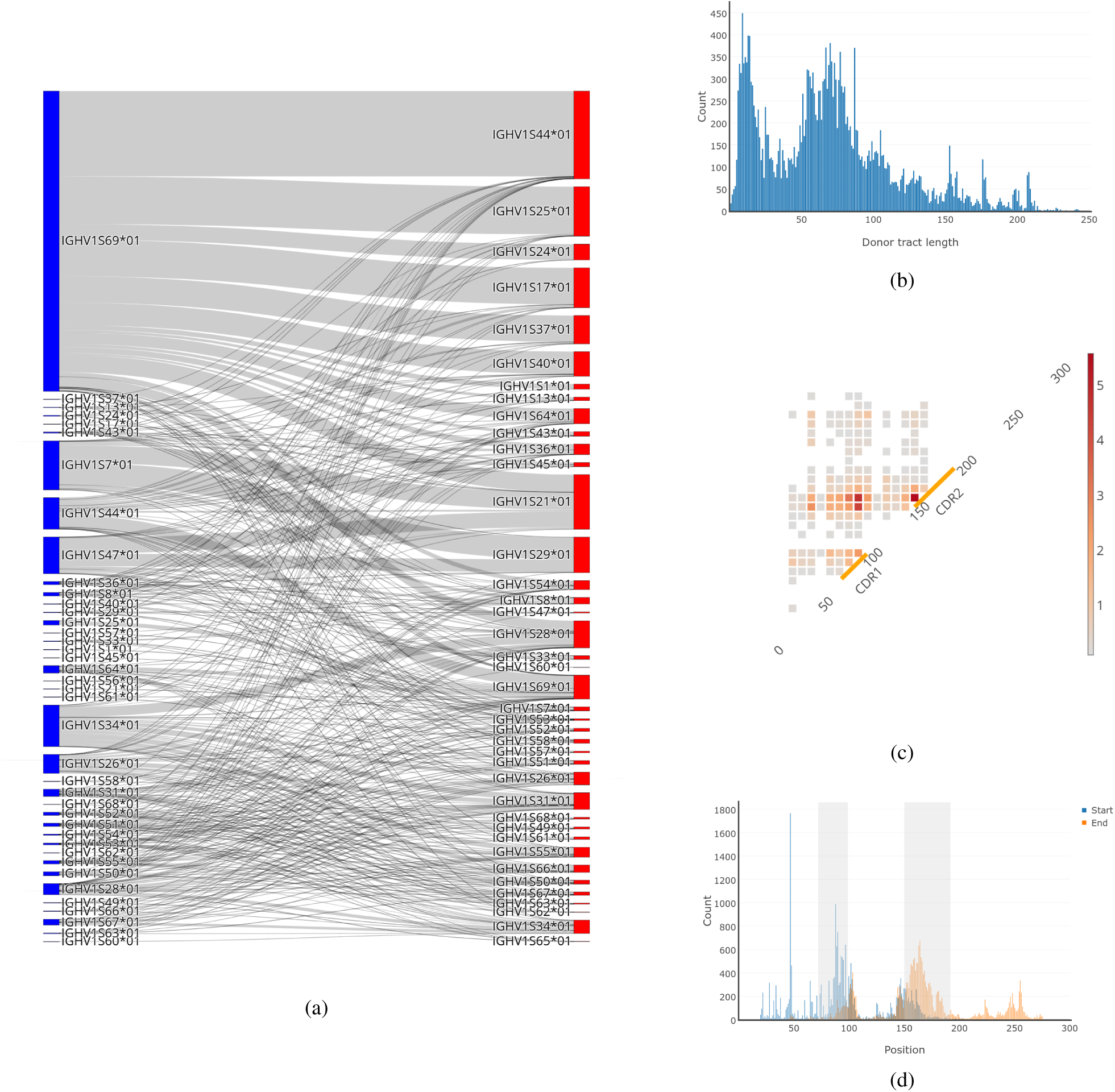
Summary of gene conversion events in immunized rabbit repertoire. a) The host V gene (left) and donor V gene (right) relationships as counts of distinct gene conversion events. b) The distribution of gene conversion tract; where the majority of gene conversion events are between 50nt-85nt. c) Donor interval map, showing start/end donor intervals in 10nt bins. Each start/end bin shows the fraction of identified donor tracts as a percentage. d) Start/end intervals of gene conversion donor tracts within their host. CDR1/2 is shaded in gray.

The start and end intervals of gene conversion events is shown in Figure 4c and 4d. There is a preference for the boundaries of donor tracts within the highlighted CDR1 and CDR2 regions. Boundaries in these regions account for 50.9% and 60.8% of the total interval boundaries for start and end, respectively. This pattern follows the model of AID mediated gene conversion; which predicts that regions of higher SHM also serve as boundaries for donor tracts more often than regions of low SHM.

## 4 Conclusion

A new and robust method for identifying gene conversion events in immunoglobulins has been presented. With `GECCO`, we are now able to characterize somatic gene conversion events in antibodies and provide a new insightful feature in the analysis of repertoires. While we applied `GECCO` only to rabbit repertoires, it can be used to discover the prevalence of gene conversion in repertoire responses in other species. Gains in recall over the only previously used approach in the literature (GENECONV and variants thereof) have been shown across a variety of donor tract lengths and mutation loads. Similarly, reduction of false positives have also been shown over non gene converted, yet mutated, simulated antibody sequences.

This new method was applied to characterizing the gene conversion events from immunized rabbit repertoires. We identified the overall frequency of gene conversion events in rabbit is ≈22% (average across three rabbits) Additionally, we found gene conversion of the dominant host gene (V1S69) showed preference for the donor gene V1S44. The two other rabbits, with different allotypes, showed different donor V gene preferences (but similar to one another). It is unclear if these variations are allotype specific, individual specific, or have an alternate explanation for their variation. Future analysis of repertoires from additional individuals, and immunzations, could shed light on any host/donor preferences.

While `GECCO` presents a new way to characterize antibody repertoires using GC-ALIGN, there are still limitations with the approach. Specifically, only a single gene conversion event can be identified. Should an antibody accept two donor tract sequences, from two different donors, currently only the single best scoring donor tract will be identified. This limitation may be reasonable in most organisms such as rabbit or human, but may be limiting in chicken where gene conversion is a main actor of immunoglobulin diversification. Additional improvements could be made to the GC-ALIGN algorithm, for example, the incorporation of a position specific mutation model would further reduce false positives, provide more accurate boundary detection of donor sites, but would require species specific training datasets. This could not only prevent false positives by adding a constraint, but it could also improve the accuracy of boundary identification of gene conversion events. By limiting the locations of gene conversion jumps between matrices, this could define where gene conversion likely occurs in regions of high homology between the donor and host sequences.

## 5 Acknowledgements

The author is grateful to Piero Bonissone for fruitful discussions throughout the development of this work and comments on the manuscript, along with thoughtful comments on the manuscript by Anand Patel, Natalie Castellana, and Thiago Lima. The author is also grateful to Thiago Lima for the preparation of rabbit sequencing libraries.

